# The MicroMap is a network visualisation resource for microbiome metabolism

**DOI:** 10.1101/2025.02.13.637616

**Authors:** Cyrille C. Thinnes, Renee Waschkowitz, Eoghan Courtney, Eoghan Culligan, Katie Fahy, Ruby A. M. Ferrazza, Ciara Ferris, Angeline Lagali, Rebecca Lane, Colm Maye, Olivia Murphy, David Noone, Saoirse Ryan, Mihaela Bet, Maria C. Corr, Hannah Cummins, David Hackett, Ellen Healy, Nina Kulczycka, Niall Lang, Luke Madden, Lynne McHugh, Ivana Pyne, Ciara Varley, Niamh Harkin, Ronan Meade, Grace O’Donnell, Bram Nap, Filippo Martinelli, Almut Heinken, Ines Thiele

## Abstract

The human microbiome plays a crucial role in metabolism and thereby influences health and disease. Constraint-based reconstruction and analysis (COBRA) has proven an attractive framework to generate mechanism-derived hypotheses along the nutrition-host-microbiome-disease axis within the computational systems biology community. Unlike for human, no large-scale visualisation resource for microbiome metabolism has been available to date. To address this gap, we created the MicroMap, a manually curated microbiome metabolic network visualisation, which captures the metabolic content of over a quarter million microbial genome-scale metabolic reconstructions. The MicroMap contains 5,064 unique reactions and 3,499 unique metabolites, including for 98 drugs. The MicroMap allows users to intuitively explore microbiome metabolism, inspect microbial metabolic capabilities, and visualise computational modelling results. Further, the MicroMap shall serve as an educational tool to make microbiome metabolism accessible to broader audiences beyond computational modellers. For example, we utilised the MicroMap to generate a comprehensive collection of 257,429 visualisations, corresponding to the entire scope of our current microbiome reconstruction resources, to enable users to visually compare and contrast the metabolic capabilities for different microbes. The MicroMap seamlessly integrates with the Virtual Metabolic Human (VMH, www.vmh.life) and the COBRA Toolbox (opencobra.github.io), and is freely accessible at the MicroMap dataverse (https://dataverse.harvard.edu/dataverse/micromap), in addition to all the generated reconstruction visualisations.

## Introduction

Human microbiome metabolism is intricately linked to health and disease.^1^ Investigating the relationships between intrinsic (e.g., genetics) and extrinsic (e.g., nutrition) contributing factors requires a holistic approach that integrates multiple layers of biomedical information along the nutrition-host-microbiome-disease axis.^2^ Computational approaches offer an attractive opportunity to tackle this dynamic and context-dependent systems biology challenge.

Genome-scale metabolic reconstructions are mathematical representations of an organism’s metabolism and enable the generation of mechanism-derived hypotheses in conjunction with the appropriate computational methods, such as constraint-based reconstruction and analysis (COBRA), *via*, e.g., the COBRA Toolbox.^3^ Ongoing efforts enabled the continuous development of the requisite human and microbiome reconstructions.^4^ Furthermore, the content of the human reconstructions Recon2^5^ and Recon3D^6^ was visualised as their metabolic network maps, ReconMaps^7^ and ReconMap3,^8^ respectively, to enable users to interactively explore human metabolism and display modelling and simulation results, such as predicted flux values using COBRA.^9^ However, no comparable microbiome network visualisation counterparts have been available thus far.

Therefore, we created the MicroMap, a manually curated network visualisation of microbiome metabolism derived from the AGORA2 resource of 7,302 human microbial strain-level metabolic reconstructions, which also captures the metabolism of 98 commonly prescription drugs.^10^ The MicroMap is also compatible with the metagenome-assembled genomes (MAG)-derived APOLLO resource of 247,092 microbial metabolicreconstructions,^11^ as all reaction and metabolite identifiers are consistent throughout both resources, and APOLLO metabolic content, i.e., number of reactions and metabolites, is smaller than of the reference-genome-derived AGORA2 reconstructions. Users may explore the standalone MicroMap to inspect the systems context of microbiome metabolism, and integrate the MicroMap with their modelling methodologies, e.g., to visualise COBRA results.

By offering an accessible and interactive means to visually explore microbiome metabolism, the MicroMap extends the capabilities of the Virtual Metabolic Human database (VMH, www.vmh.life),^8^ complementing existing visualisation tools, such as the ReconMaps for human metabolism. The MicroMap shall support research and education in human microbiome metabolism, providing critical insights into how these interactions influence health and disease.

## Methodology

### Drawing the MicroMap

The MicroMap was created within the Virtuome summer school in digital health (https://www.digitalmetabolictwin.org/virtuome) and based on the three pillars of biocomputational research, personal development, and community-engaged research. Virtuome participants were current STEM undergraduates in Ireland, including from the University of Galway, University College Cork, University College Dublin, and Trinity College Dublin. The team was dispersed across Ireland and collaborated fully online, according to best practices in systematic team empowerment.^12^

At the start, we used MATLAB to export the biochemical subsystems, unique reactions, and metabolites from the AGORA2 resource as Excel spreadsheets. We then clustered the reactions according to their assigned subsystem, which helped us to structure the rather large manual drawing task into smaller tractable segments and also provided an estimate of the required area on the map, informed by the number of reactions.

We manually drew the metabolic reactions using CellDesigner (https://celldesigner.org).^13^ Considering the large number of 8,635 AGORA2 reactions to be completed within the limited 14-week part-time Virtuome component, we decided to prioritise biotransformation reactions, and deprioritise exchange and transport reactions. We annotated the MicroMap with labels referring to biological subsystems, also including drug names, to help the user find the reactions of interest. We used colours to convey visual cohesion, e.g., by using the same colour for the same metabolites, and related colour schemes for related metabolites, such as shades of yellow for energy carriers. To facilitate efficient and consistent mapping, we established a template for recurring reaction layouts, metabolites, and associated colour schemes, which enabled us to copy and paste the ready-made constituting parts into the map in progress. After each contributor had created a handful of submaps, they integrated them into an iteratively growing, new version of the MicroMap.

Throughout, we aimed to adopt a city map-inspired design organised around subsystem nodes to facilitate intuitive navigation by the biochemically informed user. We guided the map integration process by encouraging each contributor to continuously consider the three aspects of content, design, and quality control. Content focused on the accurate representation of the parent AGORA2 reconstruction resource, design on creating a visually coherent layout, and quality control on ensuring good practice for minimising human error during the mapping process.

Once we completed a first manually assembled MicroMap draft, we used the COBRA Toolbox Metabolic Cartography functions, notably *checkCDerrors.m*, to compare the MicroMap with the AGORA2 resource, which highlighted any reaction and metabolite differences, including identifiers and reaction reversibility. We iteratively corrected the map such that all reactions and metabolites of the MicroMap are present within AGORA2. The most common mistake was the accidental introduction of a space at the beginning or end of an identifier, which are particularly challenging to spot with the naked eye. We accounted for the unique metabolite content, i.e., metabolites with the same VMH identifier without the associated compartment, by creating the *uniqueMetabolites.m* and *uniqueSpeciesInMap.m* functions.

### Visualising reconstructions

To visualise individual microbial reconstructions on the MicroMap, we created the *visualizeReconstructionsOnMap.m* COBRA Toolbox function and provided the MicroMap CellDesigner .xml file and a folder path containing the reconstruction .mat files of interest. The strain-level AGORA2 and APOLLO metabolic reconstruction files (in .mat format) were downloaded from the VMH (www.vmh.life). The AGORA2 class-, order-, family-, genus-, and species-level, and the APOLLO species-level pan reconstructions were generated using the *createPanModels.m* Microbiome Modelling Toolbox 2.0 function.^14^ For the bulk reconstruction visualisations, we used high-performance Dell PowerEdge 730 servers, with 40 CPU cores and 768Gb of RAM.

### Creating reaction presence heatmaps

To generate heatmaps of reactions present in a given set of microbial reconstruction visualisations, we created the *visualizeNormalizedRxnPresence.m* COBRA Toolbox function and provided a folder path containing the reconstruction visualisation CellDesigner .xml files of interest.

### Visualising flux vectors

We computed a flux vector using flux balance analysis (*optimizeCBmodel.m*) using the COBRA Toolbox. The flux vector was then stored in .csv format, which contained reaction identifiers in the first and associated, predicted flux values in the second column. To visualise the flux vector, we created the *addFluxFromFileWidthAndColor.m* and *visualizeFluxFromFile.m* COBRA Toolbox functions and provided the MicroMap CellDesigner .xml file and the flux vector .csv files as inputs.

### Animating flux visualisations

We obtained an example flux vector series, issued from a longitudinal series of timepoints in .xlsx format. To visualise each flux, we created the *addFluxWidthAndColor.m* and *visualizeFluxTimeseriesFromFile.m* COBRA Toolbox functions and provided the MicroMap CellDesigner .xml file and the flux vector timeseries .xlsx files as inputs.

We exported each of the resulting CellDesigner .xml files as .pdf files by opening each .xml file in CellDesigner and using ‘Export Image’. In Adobe Photoshop, we used ‘Load files into Stack’ to import each of the .pdf files into a separate layer within the same document. After cropping into an area of interest, we created a frame animation and made frames from layers. We set the frame delay to 0.25 throughout and the playback loop to ‘forever’, before exporting the resulting animation as a .gif file.

## Results

We created the MicroMap, which captures 5,064 unique reactions (representing 59% and 76% of the AGORA2^10^ and APOLLO^11^ metabolic reconstruction resources, respectively) and 3,499 unique metabolites (representing 97% and 100% of the AGORA2 and APOLLO resources respectively), including the metabolism of 98 drugs (**Figure 1a**). The city map-inspired design aims to intuitively guide the biochemically informed user, e.g., by clustering together related reactions, such as drug metabolism, and by providing 337 location signs, which indicate the relevant biochemical subsystem or drug name. Metabolites contain their VMH identifier, with the associated compartment (e.g., c for cytosol) in brackets. Reactions are labelled with their VMH identifier where feasible, i.e., not starting with a number (in which case they are displayed with their automatically assigned CellDesigner identifier). We adopted a consistent colour scheme throughout, including similar hues for related molecules, e.g., yellow for energy carriers, such as nicotinamide adenine dinucleotide (nad[c]) or adenosine triphosphate (atp[c]) (**Figure 1b**). The MicroMap CellDesigner .xml and .pdf files are freely available from the MicroMap dataverse (https://dataverse.harvard.edu/dataverse/micromap).

**Figure 1:**
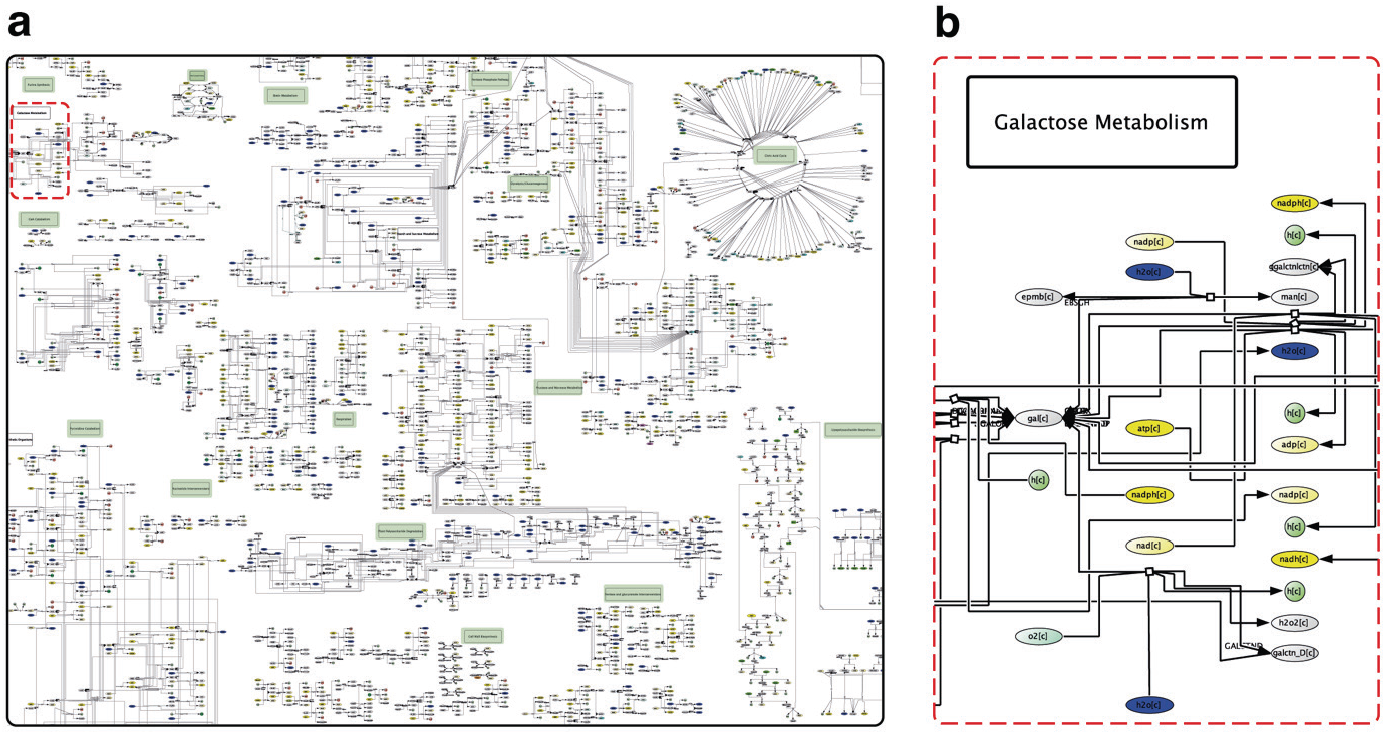
The MicroMap is a manually curated network visualisation of microbiome metabolism. **a)** A view of the MicroMap. Nodes represent metabolites and edges represent reactions. Highlighted in green are location labels, which indicate the names of metabolic subsystems or drugs to support intuitive map exploration. **b)** A magnified view of the MicroMap location within the red dashed frame of a). Metabolites contain their VMH identifier, with the associated compartment (e.g., c for cytosol) in brackets. We adopted a consistent colour scheme, e.g., by highlighting energy carriers in yellow, to support intuitive map exploration.

We then visualised microbiome reconstructions on the MicroMap to explore, compare, and contrast the known metabolic capabilities for different microbes. When comparing, e.g., the metabolic maps for Bacilli (**Figure 2a**) and Verrucomicrobia (**Figure 2b**), it is evident that Bacilli possess a relatively vast range of drug-metabolising capabilities compared to Verrucomicrobia. This visual means shall support making microbiome metabolism knowledge accessible to users, who may otherwise not be inclined to use biocomputational modelling software, e.g., in educational settings. We generated 9,742 reconstruction visualisations for AGORA2, which includes visualisations for the 7,302 strains, and for the pan-reconstructions, which include 39 classes, 78 orders, 152 families, 422 genera, and 1,749 species. We also generated 247,687 visualisations for APOLLO, including for the 247,092 strains, and for the associated 595 pan-species reconstructions. The use of two high-performance high-memory servers enabled the parallel visualisation generation to complete within 22 days – a 40-fold enhancement over the use of a conventional desktop computer, estimated to have required 2.5 years of continuous visualisation generations. Altogether, we provide the full set of 257,429 microbe visualisations in CellDesigner .xml format, freely available from the MicroMap dataverse (https://dataverse.harvard.edu/dataverse/micromap).

**Figure 2:**
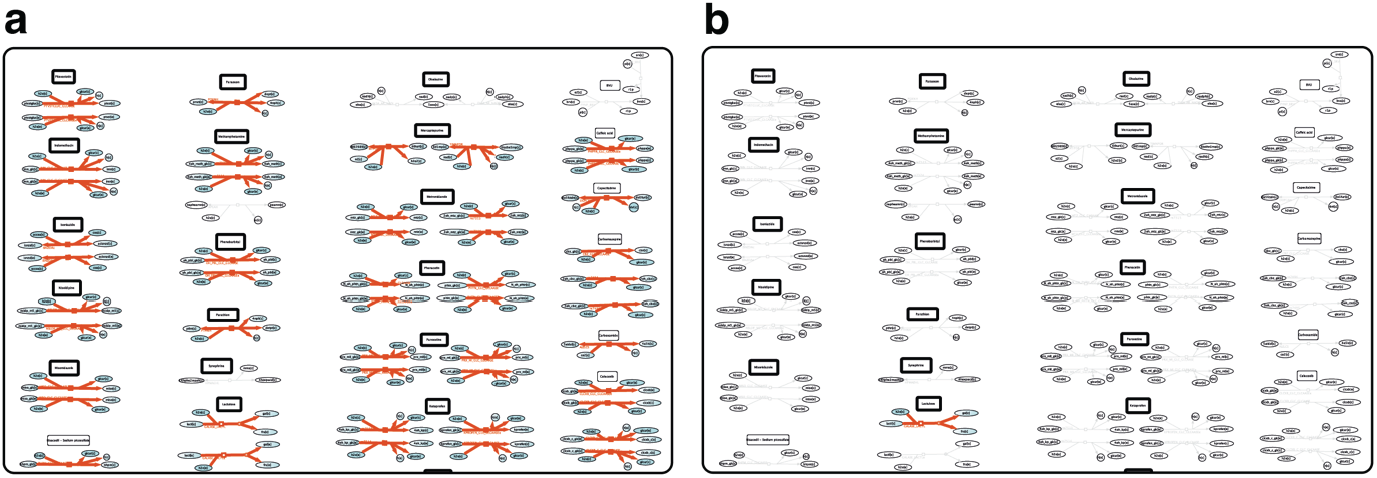
Reconstruction visualisations enable the comparison between the metabolic potential of diAerent microbes. **a)** A view of the pan-Bacilli class reconstruction visualisation (red) on the MicroMap with focus on a drug metabolism section (same as in b). **b)** A view of the pan-Verrucomicrobia class reconstruction visualisation (red) on the MicroMap with focus on a drug metabolism section (same as in a).

Further developing the capabilities for visual metabolism exploration, we implemented the capability to create heatmaps of relative reaction presence among a given set of reconstruction visualisations. We created, e.g., a heatmap, which represents the relative metabolic capabilities of 14 pseudomonas species contained within AGORA2 (**S1**). Inspection revealed that the Pseudomonas subset can, e.g., mediate some known microbiome-mediated Fluorouracil^15^ reactions, with differing metabolic capabilities among the species (**Figure 3**).

**Figure 3:**
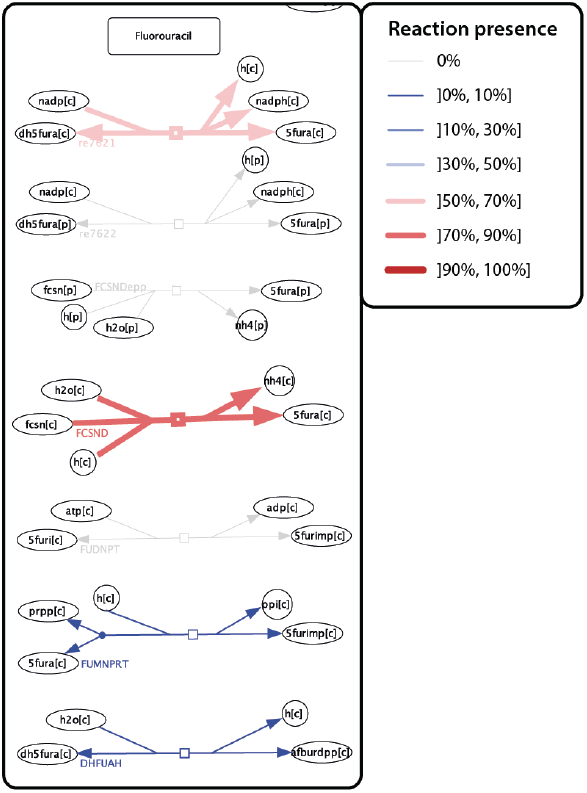
Heatmaps indicate relative reaction presence for a given set of microbial reconstructions. A view of the relative reaction presence heatmap for 14 Pseudomonas species contained within AGORA2 with a focus on Fluorouracil drug metabolism. Both hue and line width are associated with relative reaction presence, ranging from saturated blue with fine lines to saturated red with thick lines. Grey lines signify no reaction presence across the input reconstructions.

Subsequently, we used the MicroMap to visualise a flux vector which resulted from COBRA modelling and represents the flow of metabolites through a metabolic network (**Figure 4**). By extension, we implemented the bulk visualisation of flux vectors issued from a longitudinal timeseries analysis. The resulting individual maps, representing one flux vector visualisation per timepoint, offered an opportunity to create a frame-by-frame animation to highlight flux dynamics over time. The animation revealed flux changes, including in sign and magnitude, e.g., for glutamate metabolism reaction P5CD (**S2**). Consequently, such animations are a means to visually identify potential candidate pathways of interest, based on their changes over time (or lack thereof). Please note, however, that a given flux vector may be part of multiple solutions for a set of linear equations – result interpretation shall therefore be informed by the specific COBRA methodology used.

**Figure 4:**
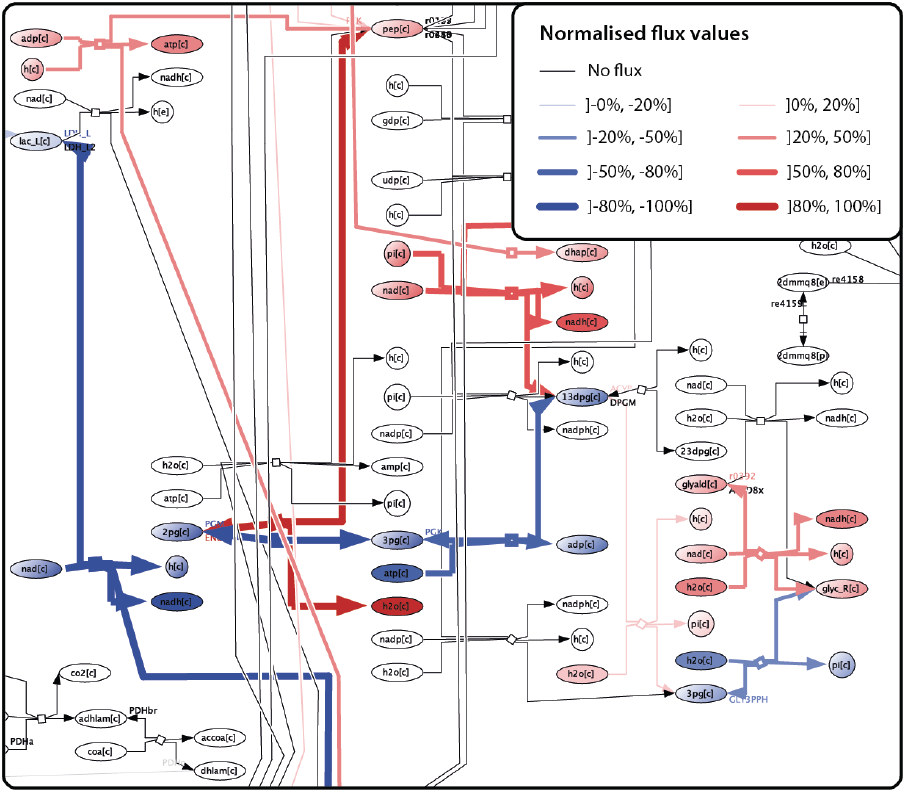
COBRA modelling results can be visualised on the MicroMap. A view of an example flux vector visualisation on the MicroMap. Blue indicates negative flux values, red indicates positive flux values. Line width and hue reflect the magnitude of the reaction-associated flux. Grey lines signify that no flux is associated with the reaction.

## Discussion

The MicroMap is a manually curated metabolic network visualisation of the human microbiome. The MicroMap captures the metabolic content of over a quarter million microbial genome-scale metabolic reconstructions, encompassing ∼5,000 unique reactions and ∼3,500 unique metabolites, including from 98 drugs. The MicroMap serves to 1. visually explore microbiome metabolism, 2. compare and contrast the metabolic capabilities of different microbes, and 3. visualise computational modelling results.

The MicroMap is fully integrated within the VMH and therefore, seamlessly interconnects with all VMH resources, including human metabolism. The MicroMap complements the portfolio of biocomputational tools for studying human-microbiome interactions, such as the nutrition-gut-brain-health axis. The implemented visualisation methodologies, i.e., COBRA Toolbox functions, are transferable to related contexts, such as human metabolism-related visualisations involving the ReconMap3. Likewise, the already existing Metabolic Cartography functions are compatible with microbiome visualisations involving the MicroMap.

Beyond its applications in research and development, the MicroMap also constitutes an educational tool for exploring, learning about, and communicating aspects on microbiome metabolism. The visualisations aim to be accessible to users beyond computational metabolic modelling, including the wider scientific communities, and stakeholders with a general interest in metabolism. After dabbling in generating heatmaps and animations, we encourage the user community to explore and propose new creative use cases for iterative improvement not only of the MicroMap, but also metabolic modelling visualisations in general.

## Data availability

The MicroMap and all the created AGORA2 and APOLLO reconstruction visualisations are freely accessible and can be downloaded from the MicroMap dataverse (https://dataverse.harvard.edu/dataverse/micromap). The CellDesigner .xml files can be viewed, explored, and further edited using the open-access CellDesigner software (https://celldesigner.org). The COBRA Toolbox functions are available from within the COBRA Toolbox (opencobra.github.io).

## Acknowledgements

This work has been funded by the European Research Council (ERC) under the European Union’s Horizon 2020 research and innovation programme (grants #757922 & 101125633), the European Union’s Horizon 2020 research and innovation programme under the Marie Skłodowska-Curie Actions (grant #859890), the National Institute on Aging (grants #1RF1AG058942 & 1U19AG063744), and the Science Foundation Ireland (grant #12/RC/2273-P2). We thank Anna Sheehy and Tim Hensen for providing flux vector examples.

## Notes

https://dataverse.harvard.edu/dataverse/micromap

### Competing Interest Statement

The authors have declared no competing interest.

