## Supplementary figures and images for "The MicroMap is a network visualisation resource for microbiome metabolism"

### S1

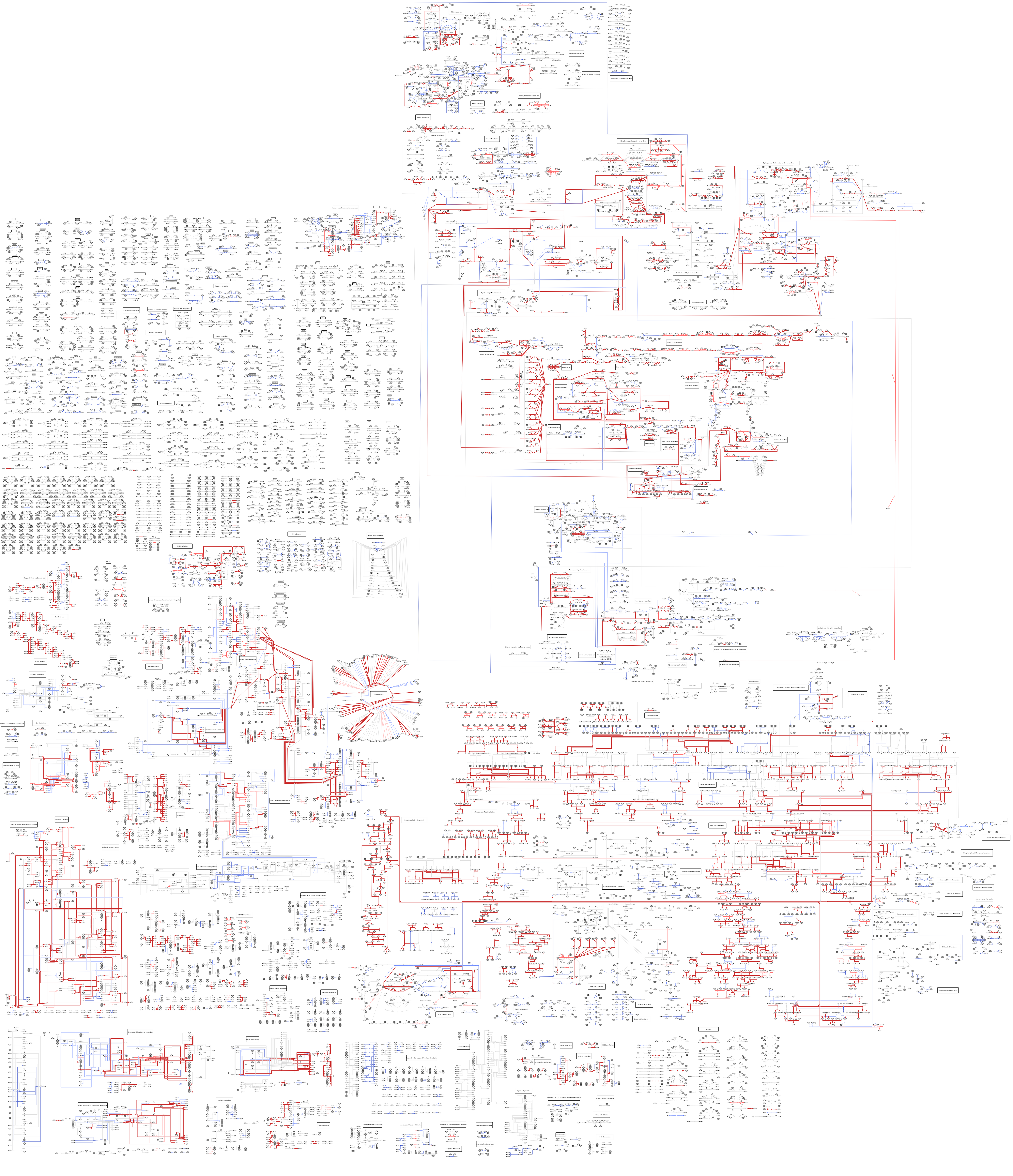

### S2

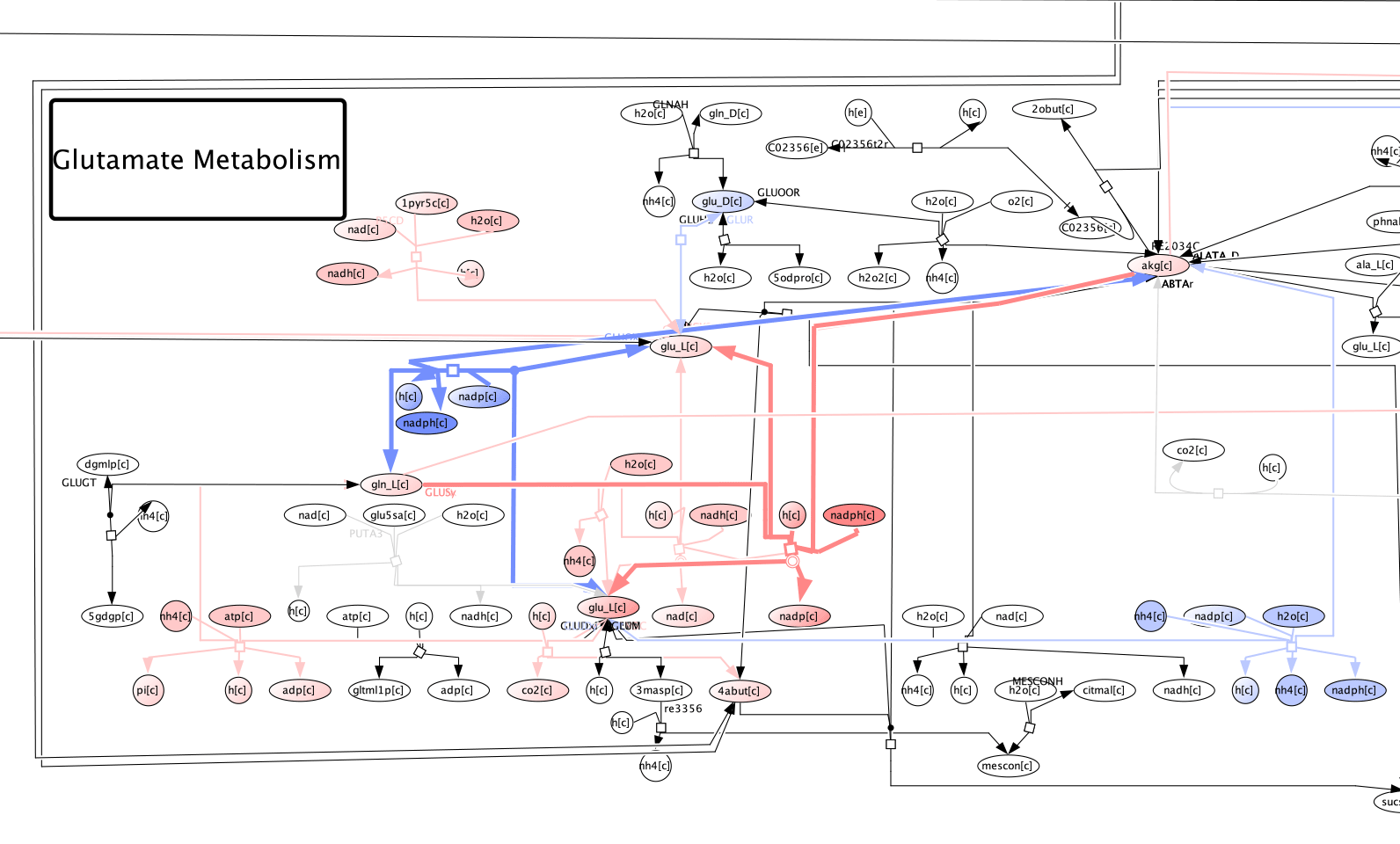
